# The maternal environment interacts with genetic variation in regulating seed dormancy in *Arabidopsis thaliana*

**DOI:** 10.1101/117879

**Authors:** Envel Kerdaffrec, Magnus Nordborg

## Abstract

Seed dormancy is a complex adaptive trait that controls the timing of seed germination, one of the major fitness components in many plant species. Despite being highly heritable, seed dormancy is extremely plastic and influenced by a wide range of environmental cues. Here, using a set of 92 *Arabidopsis thaliana* lines from Sweden, we investigate the effect of seed maturation temperature on dormancy variation at the population level. The response to temperature differs dramatically between lines, demonstrating that genotype and the maternal environment interact in controlling the trait. By performing a genome-wide association study (GWAS), we identified several candidate genes that could account for this plasticity, two of which are involved in the photoinduction of germination. Altogether, our results provide insight into both the molecular mechanisms and the evolution of dormancy plasticity, and can serve to improve our understanding of environmentally dependent life-history transitions.

**Highlight:** The effect of low seed-maturation temperatures on seed dormancy is highly variable in *Arabidopsis thaliana* accessions from Sweden, denoting strong genotype-environment interactions, and a genome-wide association study identified compelling candidates that could account for this plasticity.

## Introduction

Life-stage transitions, the timing of which is critical to plant fitness, are regulated by both genes and the environment, usually in interaction (G × E) (El-Soda *et al.*, 2014; Koornneef *et al.*, 1998; Chiang *et al.*, 2013). Plants have evolved ways to sense and integrate environmental inputs in order to adjust their life cycle to seasonal environments, the best-known example of which is vernalization and the perception of winter cold (Amasino, 2004). In *A. thaliana,* vernalization results in the stable repression of the central regulator *FLOWERING LOCUS C* (*FLC*), a prerequisite for the vegetative-to-reproductive transition to occur (Sheldon *et al.*, 2000).

While flowering time determines the reproductive environment, seed dormancy, another major life-history trait crucial for local adaptation, regulates the timing of germination and determines the post-germination environment (Donohue, 2002; Donohue *et al.*, 2010; Chiang *et al.*, 2013). Dormancy is highly plastic and can be modulated both by pre-and post-dispersal environmental cues such temperature, light, and to a lesser extent, nitrate (Fenner, 1991; Baskin and Baskin, 1998; Footitt *et al.*, 2013; Huang *et al.*, 2015; Penfield and MacGregor, 2016; Finch-Savage and Footitt, 2017). In particular, low temperatures during seed production dramatically increase dormancy in *A. thaliana* (Chiang *et al.*, 2011; Kendall *et al.*, 2011; Huang *et al.*, 2014; He *et al.*, 2014) as well as in other species such as wheat (Reddy *et al.*, 1985) and wild oat (Peters, 1982).

Central to this temperature-dependent process in *A. thaliana* is the upregulation of a major genetic determinant of seed dormancy variation, *DELAY OF GERMINATION 1* (*DOG1*) (Bentsink *et al.*, 2006; Chiang *et al.*, 2011; Kendall *et al.*, 2011). The *DOG1* locus exhibits genetic signatures suggestive of local adaptation, and field experiments have emphasized its pivotal role in controlling the timing of germination in wild *A. thaliana* populations (Huang *et al.*, 2010; Kronholm *et al.*, 2012; Postma and Ågren, 2016; Kerdaffrec *et al.*, 2016). Independently of their action on *DOG1,* low seed maturation temperatures induce deep dormancy by increasing the abscisic acid (ABA) / gibberellins (GA) ratio, two antagonistic phytohormones repressing and activating germination, respectively (Chiang *et al.*, 2011; Kendall *et al.*, 2011). Finally, low temperatures can promote coat-imposed dormancy by altering seed coat permeability through the upregulation of the flavonoids biosynthesis pathway, both during seed production (MacGregor *et al.*, 2015) and vegetative phase (Chen *et al.*, 2014).

Thus, it is clear that the induction of primary dormancy is regulated by both genetic and environmental factors, and the fact that distinct genotypes differ in their response to low temperatures indicates that genotype-environment interactions partly control the trait (Schmuths *et al.*, 2006; Penfield and Springthorpe, 2012; He *et al.*, 2014; Burghardt *et al.*, 2016a). A direct consequence of this plasticity is that similar germination trajectories, defined as the evolution of the germination phenotype over time, can be promoted by different combinations of genotypes and environments. For example, strong *DOG1* alleles combined with a warm maternal environment and weak *DOG1* alleles combined with a cold maternal environment can both lead to highly dormant phenotypes (Burghardt *et al.*, 2016a).

Field studies have demonstrated that the maternal environment contributes greatly to seed dormancy variation under natural conditions (Postma and Ågren, 2015). Thus, given the existence of strong selection for timing of germination (Donohue *et al.*, 2005; Huang *et al.*, 2010; Postma and Ågren, 2016; Kerdaffrec *et al.*, 2016), it has been speculated that the temperature-dependent regulation of primary dormancy is adaptive. For example, it may provide the mother plant with information regarding the seasonal environment, information that can be used to set dormancy appropriately (Galloway and Etterson, 2007). In addition, this mechanism is expected to enable bet-hedging strategies, in which seeds from the same population — or plant — express various dormancy phenotypes and germinate throughout the year to maximize fitness in unpredictable environments (Venable and Brown, 1988; Simons and Johnston, 2006; Wilczek *et al.*, 2010; Mitchell *et al.*, 2016).

Although genotype-environment interactions have previously been reported to influence dormancy and germination plasticity in *A. thaliana* (Schmuths et al., 2006; Penfield and Springthorpe, 2012; He et al., 2014; Burghardt et al., 2016a), the extent of this phenomenon and whether it is universal at the species level is unknown. Moreover, the genetic basis of the differential response to seed maturation temperatures, and more generally, of G × E variation, remains to be thoroughly investigated. Here, by growing a set of *A. thaliana* lines from Sweden in two different environments, we assess the effect of temperature on seed dormancy in a local sample, before performing a GWAS to identify the genes responsible for the observed variation.

## Material and methods

### Plant material and phenotyping

The 92 Swedish lines used in this study (Table S1) were previously described by Kerdaffrec *et al.* (2016). For each genotype, six biological replicates were vernalized for eight weeks (4°C, standard long days, 90% humidity). Then, three randomly chosen replicates were placed in a warm environment (21°C day, 16°C night), while the other three received a cold treatment (15°C day, 10°C night). Both treatments were applied from rosette stage to ripening and seed harvest. Seeds were harvested when about 50% of the siliques of a given plant had come to maturity and were subsequently placed in dry environment for after-ripening (30% relative humidity, 16°C, dark). The germination rate seeds after-ripened for 21, 63 and 105 days (GR21, GR63 and GR105) was estimated by scoring radicle emergence after a week of incubation at 23°C under standard long days (Alonso-Blanco *et al.*, 2003; Kerdaffrec *et al.*, 2016).

### Broad sense heritability

Broad sense heritabilities (H) were calculated as the genotypic variance divided by the total variance. Both variances were estimated using a linear mixed model from the lme4 package in the R framework (R Core Team, 2014). The model was as follows:

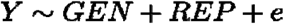

where *GEN* is the genotype (line), *REP* is technical replicate, and *e* is the error. All variables were fitted as random effects.

### Variance components analysis

The variance-component analysis was described earlier by Sasaki *et al.* (2015). It was carried out in LIMIX (Lippert *et al,* 2014) using the following model:

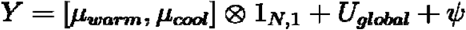

where *μ_warm_* and *μ_cool_* are environment specific mean values, *U_global_* is a matrix of global relatedness fitted as random effect, and *ψ* is noise.

### Genome-wide association mapping

Genome-wide scans were performed on arcsine transformed mean phenotypic values using a mixed-model accounting for population structure and SNPs derived from the 1001 genomes project (Kang *et al.*, 2010; Zhang *et al.*, 2010; The 1001 Genomes Consortium, 2016). The analysis was performed using the GWA-Portal (https://gwas.gmi.oeaw.ac.at/; Seren *et al.*, 2012), and both settings and results can be viewed interactively online: http://bit.ly/2lkzAsp. In this manuscript, rare alleles (minor allele frequency < 14%) were filtered and a 5% genome-wide significance threshold was determined using Bonferroni correction. A given peak was considered to have a ‘specific’ effect when its highest score for any of the three warm phenotypes did not exceed 2, or a ‘common’ effect when its score for at least one phenotype in both environment was higher than 4. Peaks that did not meet any of these arbitrary requirements were considered to have an ‘unclear’ effect.

### Gene enrichment analyses

The seed dormancy *a priori* candidate gene list (91 genes; Table S2) was built regardless of the GWAS results by searching the literature (Kendall *et al.*, 2011; Graeber *et al.*, 2012) and the ARAPORT11 database. In a nutshell, a gene was considered significantly associated when at least one SNP in the 40 kb surrounding its coding sequence had a *P* value lower than 10^−4^. Lists of non *a priori* significantly associated TAIR11 genes were built in a similar manner for each phenotype, and the overrepresentation of *a priori* genes in the resulting lists was assessed using Fisher exact tests.

### Data availability

Both raw and processed data used in this manuscript have been uploaded at GitHub: https://github.com/Gregor-Mendel-Institute/dormancy.

## Results

### Seed dormancy variation in the Swedish population

We have previously shown that there is extensive natural variation for seed dormancy among *A. thaliana* lines from Sweden, and that focusing on such a local sample increases one’s ability to detect local adaptation (Kerdaffrec *et al.*, 2016). For these reasons, we used a set of 92 Swedish lines (Table S1) to assess the effect of the maternal environment on dormancy variation. Several replicates of each line were vernalized for 8 weeks before being allowed to self-fertilize and produce seed under either warm (21°C) or cold (15°C) temperatures. The dormancy levels of the seeds were estimated by measuring their germination rate (GR) as a function of seed age (21, 63 and 105 days), also known as after-ripening, resulting in six different dormancy primary phenotypes (GR21, GR63 and GR105, for both warm and cold treatments) (Fig. 1).

**Figure 1.**
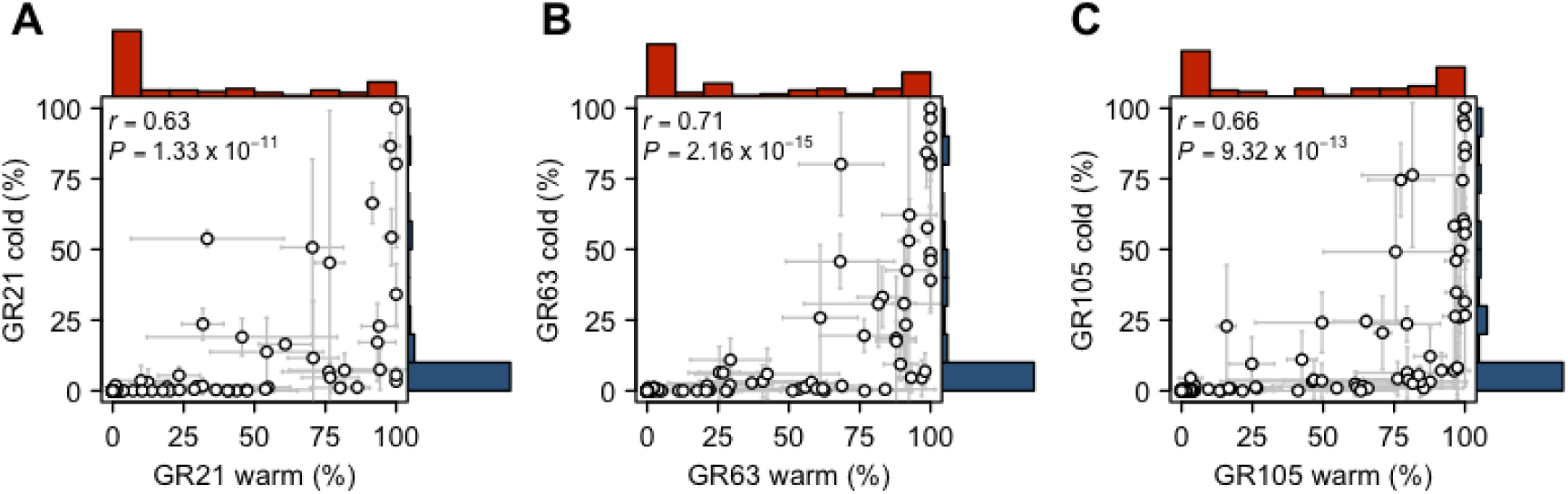
The effect of low seed maturation temperatures on seed dormancy variation. Scatter plots and histograms showing the relationship between dormancy traits as well as their phenotypic distribution. Seeds were produced either under warm (21°C; red) or cold (15°C; blue) conditions and after-ripened either for (A) 21, (B) 63 or (C) 105 days. Error bars represent the standard deviation within genotypes (n = 3, in few cases n = 2).

As expected, genotypes that experienced warm maternal temperature displayed great variation for seed dormancy, although more than one third of the lines remained dormant even after 105 days (Fig. 1). In contrast, the vast majority of seeds produced at cold maternal temperature remained dormant throughout the experiment, in line with previous studies that have shown that a decrease in seed maturation temperature generally induces a deeper dormancy (Chiang *et al.*, 2011; Kendall *et al.*, 2011; Penfield and Springthorpe, 2012; Huang *et al.*, 2014; He *et al.*, 2014; Burghardt *et al.*, 2016a).

All six phenotypes were correlated, especially within temperature treatments (Fig. S1), and broad sense heritabilities were remarkably high (ranging from 0.85 to 0.94), although they became moderate when considering lines with intermediate phenotypes only (GR > 5% and < 95%; ranging from 0.67 to 0.75; Table 1). To assess the relative effects of genes and the environment on dormancy variation, we performed a variance components analysis, in which we modelled the effect of genotype (G; line), environment (E; maturation temperature) and the interaction of both (G × E; line x maturation temperature) (Sasaki *et al.*, 2015). G and G × E effects contributed equally after 21 days (37 and 38 percent, respectively), but the purely genetic effect increased over time. Environment effects were responsible for 20% of the variance regardless of time point (Table 2). Although the accuracy of this analysis is limited because of the relatively small sample size (due to many of the tested lines being too dormant to be ‘informative’), these findings agree with previous studies and indicate that the dormancy variation observed in the Swedish sample is, to a large extent, explained by G × E effects (Schmuths *et al.*, 2006; Penfield and Springthorpe, 2012; He *et al.*, 2014; Burghardt *et al.*, 2016a).

**Table 1.** Heritability of seed dormancy traits. Broad sense heritabilities (H) were calculated using either the full sample or subsets of lines (whose number is indicated by n) with intermediate phenotypes.

|  | H<sub>warm</sub> | n<sub>warm</sub> | H<sub>cold</sub> | n<sub>cold</sub> |
| --- | --- | --- | --- | --- |
| Full sample |  |  |  |  |
| GR21 | 0.93 | 92 | 0.85 | 92 |
| GR63 | 0.93 | 92 | 0.87 | 92 |
| GR105 | 0.94 | 92 | 0.87 | 92 |
| Intermediate lines (GR ≥ 5% and GR ≤ 95%) |  |  |  |  |
| GR21 | 0.75 | 38 | 0.65 | 20 |
| GR63 | 0.75 | 42 | 0.65 | 27 |
| GR105 | 0.71 | 38 | 0.67 | 32 |

**Table 2.** Genetic and environmental effects on seed dormancy variation.

|  | G | E | G x E | noise |
| --- | --- | --- | --- | --- |
| GR21 | 37.27 | 20.48 | 38.13 | 4.13 |
| GR63 | 50.06 | 19.46 | 19.18 | 11.30 |
| GR105 | 47.75 | 22.33 | 24.86 | 35.06 |

### Genetic variation in the response to low seed maturation temperatures

It is thus clear that the effect of the maternal environment differs between Swedish genotypes. About one third of the mild- and non-dormant lines appeared to be relatively insensitive to the maternal environment after 21 days, and one line, Gro-3, reached 100% of germination in both conditions. In sharp contrast, other non-dormant lines such as Löv-1 and T480 were heavily affected by the maternal environment and displayed very low germination rates when seeds were produced at low temperatures, even after 105 days of after-ripening (Fig. S2).

To characterize the genetic variation in the response to low seed maturation temperatures, we clustered lines based on their germination phenotypes across environments and time. We identified six main clusters representing distinct germination behaviors (Fig. 2). The largest cluster (cluster 1; n = 43) is not only deeply dormant but also insensitive to after-ripening, making it ‘non-informative’ in the sense that it is not possible to assess its degree of responsiveness to temperature. Two smaller clusters 2 (clusters 2; n = 12, and 3; n = 14) display shallow to mild dormancy at 21°C that can be lifted with after-ripening, but the cold treatment induces deep dormancy that cannot be broken. Two clusters (5 and 6, n = 11 and n = 8, respectively) are both non-dormant at 21°C, but cold seed maturation temperatures dramatically increased dormancy levels of one, but had very little effect on the latter. This last observation clearly confirms that there is natural genetic variation in the response to low temperatures and that genotype-environment interactions underlie dormancy variation in the Swedish population. A spectacular example of this differential response can be found in the opposite trajectories of Gro-3 (cluster 6) and Löv-1 (cluster 5) (Fig. 2 and S2). Finally, a small cluster (cluster 4; n = 4) shows low dormancy regardless of the maternal environment, demonstrating that the degree of responsiveness is independent of the dormancy level.

**Figure 2.**
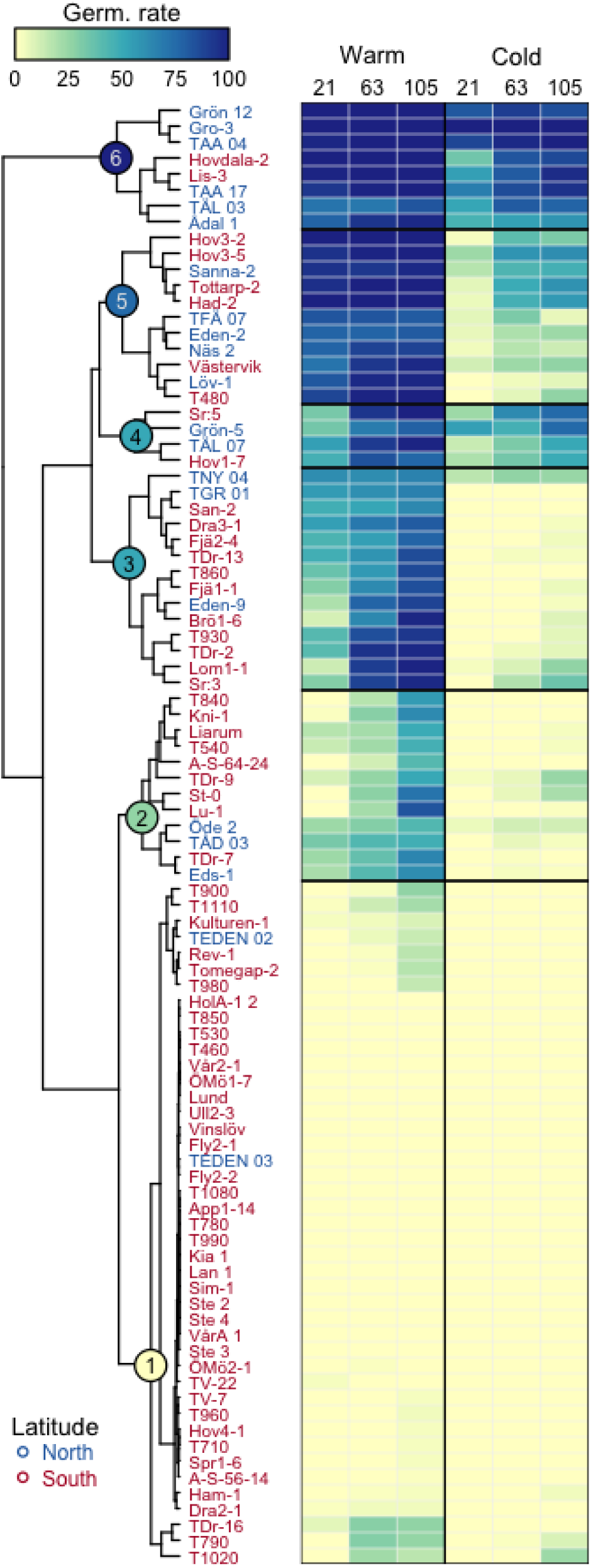
The effect of maternal environment on germination phenotypes and trajectories. Clustering dendrogram reporting the high disparity in germination trajectories across maternal environments (warm or cold) and time (21, 63 or 105 days of after-ripening). The six major clusters are numbered from 1 to 6 and are indicated with colored circles on the nodes of the dendrogram. Lines names are colored according to latitude of origin: south Sweden (red) is defined as the region below 60°N and north Sweden (blue) as the region above 60°N. Heatmap colors represent germination phenotypes, with darker shades indicating higher germination rates.

Burghardt *et al.* (2016a) have previously shown that similar germination phenotypes and trajectories can be reached via different paths, and although we only assess the effect of pre-dispersal temperatures, our results go in the same direction. For instance, cluster 5 and to some extent cluster 3 are non-dormant when seeds are produced at 21C, but lower seed maturation temperatures induced a deep dormancy, comparable to that of cluster 1.

### Geographic pattern of the response to low seed maturation temperatures

It is well established that seed dormancy correlates with latitude and climate variables such as temperature and precipitation, with northern lines generally being less dormant than southern ones (Kronholm *et al.*, 2012; Debieu *et al.*, 2013; Kerdaffrec *et al.*, 2016). This geographic pattern, thought to reflect local adaptation, was also observed in this study: both GR21 warm (Fig. 3A; *r* = 0.5, *P* = 4.44 × 10^−7^) and cold (Fig. 3B; *r* = 0.5, *P* = 4.61 × 10^−7^) are correlated with latitude. However, we note that these relationships are not strict, possibly reflecting adaptation to microenvironmental variation.

**Figure 3.**
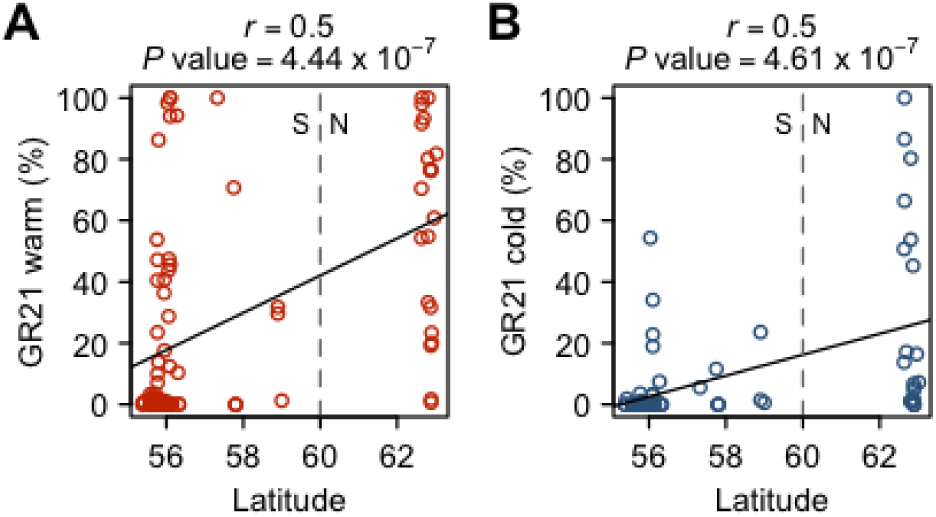
The geographic pattern of the dormancy variation. Correlation between latitude and either (A) GR21 warm or (B) GR21 cold. As in Fig. 2, We define south Sweden (S) as the region below 60°N and north Sweden (N) as the region above 60°N. See Fig. S1 for the correlations between latitude and the other phenotypes.

When grown under cold maternal conditions, almost all non-dormant lines from the south appear to be severely affected, and exhibit strongly reduced germination rates. Northern lines, however, display a greater variation in their response to low seed maturation temperature, with some genotypes being insensitive (Fig. 3B). This suggests that the response not only varies along a latitudinal gradient but also at a very local scale. These findings are nicely captured by the above-mentioned clustering approach, in which most lines from northern Sweden are binned in the sensitive and insensitive clusters 5 and 6, respectively (Fig. 2).

### GWAS for the response to low seed maturation temperatures

To uncover the polymorphisms underlying the differential response to low seed maturation temperatures, we assessed the significance of associations between the seed dormancy phenotypes and genome-wide SNP markers from the 1001 genomes project using a mixed-model accounting for population structure (Kang *et al.*, 2010; Zhang *et al.*, 2010; The 1001 Genomes Consortium, 2016). Four lines with missing genotype information were removed from the dataset, bringing the number of lines to 88 (Table S1). GWAS results were very comparable between time points, as expected given the strong correlations between traits within treatments (Fig. S1), but they differed markedly between treatments, with no strong association for the warm phenotypes while several peaks reached genome-wide significance for the cold phenotypes (Fig. 4).

**Figure 4.**
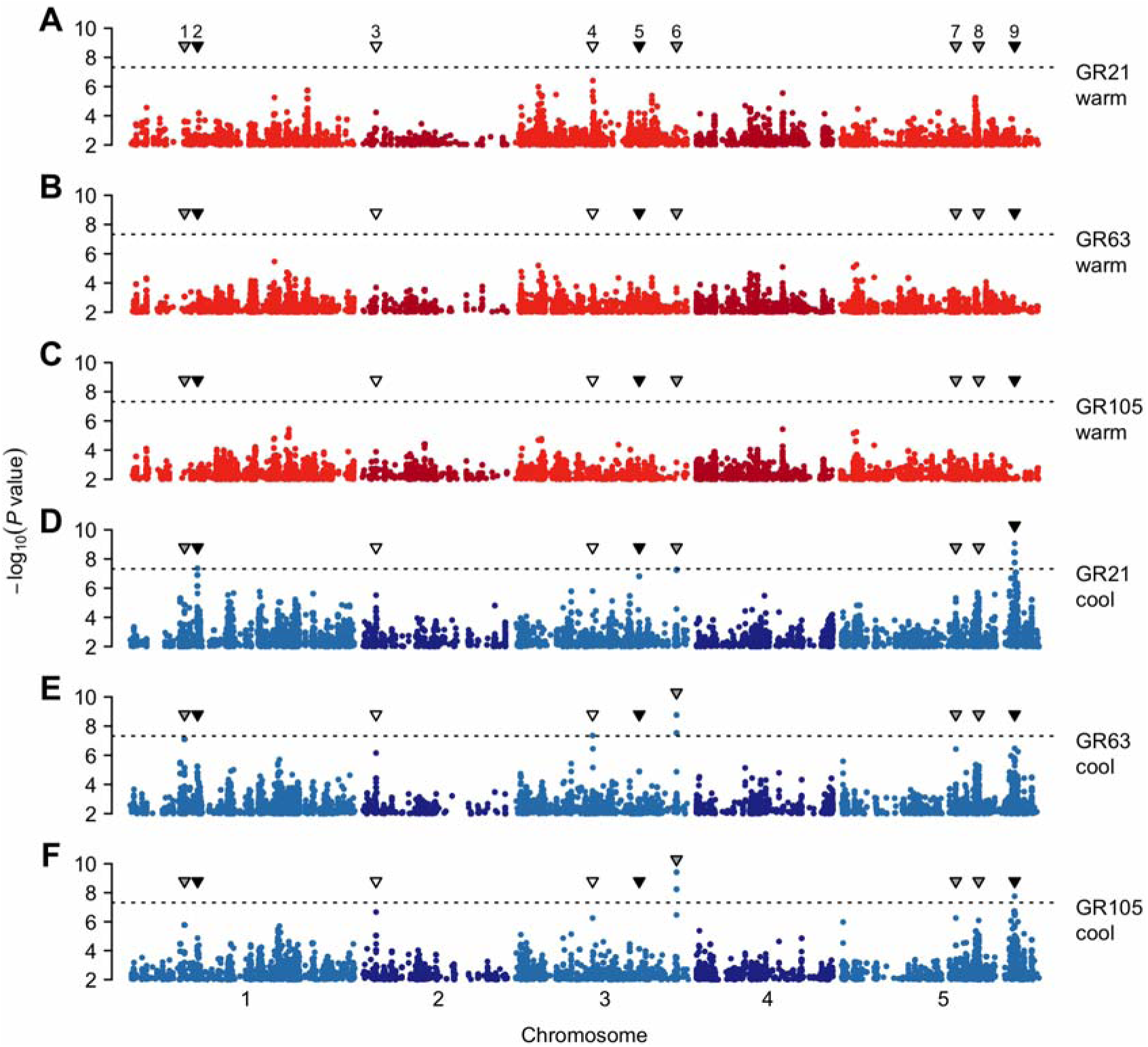
GWAS for seed dormancy phenotypes. Manhattan plots of genome-wide association results for germination rate of seeds set either in (A-C) warm or (D-F) cold environments and after-ripened for (A and D) 21, (B and E) 63 or (C and F) 105 days. The dotted horizontal line indicates a significance level of 0.05 after Bonferroni correction for multiple testing. Triangles show the position of the nine peaks with *P* values lower than 10^−6^ for at least one phenotype. Triangle color indicates the type of effect: white, ‘common’; black, ‘specific’; grey, ‘unclear’. Are only displayed SNPs with a minor allele frequency greater than or equal to 14%. The GWAS results can be viewed interactively online: http://bit.ly/2lkzAsp

To explore the GWAS results, we first performed an *a priori* gene enrichment analysis using a set of 91 genes with known or predicted function in seed dormancy regulation (Table S2). No enrichment was detected for any of the six traits, but the fact that *DOG1* is the most strongly associated *a priori* candidate suggests that some of the associations are true signals rather than noise (Table S3).

Next, we looked at the associations in greater detail, limiting ourselves to an arbitrarily chosen *P* value cutoff of 10^−6^. This yielded a total of nine regions across the six phenotypes, regions that we classified into three categories (see Materials and methods): those with a ‘common’ effect (they tend to have a similar effect on both warm and cold phenotypes), those with a ‘specific’ effect (they tend to influence only the cold phenotypes), and last, those with an ‘unclear’ effect (Fig. 4 and Table 3).

**Table 3.** Summary of the GWAS for seed dormancy phenotypes. Are listed top SNPs for the nine GWAS peaks with a score [-log_10_(*P* value)] greater than or equal to six for at least one phenotype. Genome-wide significant scores (>= 7.33) are highlighted in bold. MAF stands for minor allele frequency.

| Peak | Chr. | Pos. | MAF | Warm |  |  | Cold |  |  | Effect | Candidate genes |
| --- | --- | --- | --- | --- | --- | --- | --- | --- | --- | --- | --- |
|  |  |  |  | GR21 | GR63 | GR105 | GR21 | GR63 | GR105 |  |  |
| 1 | 1 | 7,381,921 | 0.28 | 3.60 | 3.06 | 2.47 | 4.86 | 7.10 | 5.77 | unclear | *URGT2* |
| 2 | 1 | 9,148,998 | 0.14 | 0.33 | 0.47 | 0.46 | **7.36** | 5.21 | 4.86 | specific | *SNS1* |
| 3 | 2 | 1,882,558 | 0.26 | 4.23 | 3.69 | 3.89 | 5.52 | 6.16 | 6.67 | common | - |
| 4 | 3 | 10,504,938 | 0.22 | 5.32 | 3.79 | 2.57 | 5.80 | **7.35** | 6.26 | common | - |
| 5 | 3 | 16,820,806 | 0.14 | 1.97 | 1.83 | 1.39 | 6.82 | 4.89 | 4.11 | specific | *PHOT1* |
| 6 | 3 | 21,867,111 | 0.15 | 3.19 | 3.36 | 2.39 | 7.24 | **8.76** | **9.42** | unclear | - |
| 7 | 5 | 15,630,623 | 0.14 | 2.80 | 3.27 | 2.88 | 5.32 | 6.42 | 6.26 | unclear | - |
| 8 | 5 | 18,726,653 | 0.20 | 2.96 | 2.06 | 1.58 | 4.99 | 5.21 | 6.09 | unclear | *DOG1* |
| 9 | 5 | 23,596,831 | 0.14 | 1.22 | 1.80 | 1.75 | **9.07** | 6.48 | **7.76** | specific | *PHOT2, SIP1* |

The only two associations with a ‘common’ effect, peaks 3 and 4, lie on chromosomes 1 and 2, respectively, but no clear candidate could be spotted in the vicinity of those peaks. The major ‘specific’ hit on chr. 5 (peak 9) falls directly in *SOS3-INTERACTING PROTEIN 1* (*SIP1*), a gene encoding a SnRK3-type protein kinase likely to be involved in stress and ABA signaling (Halfter *et al.*, 2000; Hrabak *et al.*, 2003). However, as the peak is quite broad (more than 100 kb; see Fig. S3), we also examined other genes in the region. Among those was *PHOTOTROPIN2* (*PHOT2*), a promising candidate not only because of its role in the photoregulation of germination, but also because *PHOT1,* a gene with similar function, was identified on chr. 3 below peak 5 (also ‘specific’). Both genes encode blue/UV-A photoreceptors and are known to be involved in the transition from dormancy to germination (Jedynak *et al.*, 2013). The third ‘specific’ association, peak 1, colocalizes with *SnRK2-substrate 1* (*SNS1*), a gene required in ABA signaling (Umezawa *et al.*, 2013). Among the four remaining peaks with an ‘unclear’ effect, we note the presence of *URGT2,* a seed specific gene controlling mucilage formation (chr. 1, peak 2) (Rautengarten *et al.*, 2014). Finally, in contrast with our previous work (Kerdaffrec *et al.*, 2016), no strong association was detected at the *DOG1* locus (peak 8, see Fig. S4), a point we discuss below.

## Discussion

The regulation of seed dormancy by maternal environment temperature has been described in numerous plant species and appears to be conserved among higher plants (Penfield and MacGregor, 2016). In *A. thaliana,* the underlying mechanism has mainly been studied at the molecular level, using very specific, often artificial backgrounds (He *et al.*, 2014; Burghardt *et al.*, 2016a). Few studies have approached this temperature-dependent regulation from a natural variation perspective, and both its extent and genetic basis remain unknown. Here, by focusing on a set of Swedish lines, we aimed to characterize this phenomenon at the population level and to identify its underlying genetic basis.

### The role of maternally-regulated dormancy in plant adaptation

Despite the prevailing deep dormancy in the Swedish sample, several lines let us assess the effect of maternal temperature on seed dormancy variation (Fig. 1). In agreement with previous reports (Penfield and Springthorpe, 2012; He *et al.*, 2014; Burghardt *et al.*, 2016a), we find that, although low seed production temperatures generally increase primary dormancy, the effect differs between lines, indicating that the trait is influenced by genotype-environment interactions. This observation is further supported by a variance component analysis, which estimates that almost 40% of the dormancy variation in the Swedish sample is due to G × E effects (Table 2; GR21). This, and the fact that high G × E variation was previously observed in a set of world-wide lines (Penfield and Springthorpe, 2012), suggests that the maternal regulation of seed dormancy by environmental cues is conserved not only at the population, but also at the species level.

By combining different genotypes and seed maturation temperatures, Burghardt *et al.* (2016a) have demonstrated that identical germination trajectories can be achieved by going down different paths. Likewise, we found that cold seed maturation temperatures can produce highly dormant phenotypes, similar in depth to those caused by genetic effects, showing that environmental variation can have large repercussions on the expression of genetic variation (Fig. 2).

Because the timing of germination is one of the major fitness components for *A. thaliana* (Postma and Ågren, 2016; Kerdaffrec *et al.*, 2016), the idea that such maternal regulation may be adaptive is attractive, although the rationale for its existence in this species is yet to be established (Burghardt *et al.*, 2016a). In Sweden, where *A. thaliana* mainly behaves as a winter annual, seed dispersal usually occurs in spring, and germination in fall. Therefore, we hypothesize that Swedish populations use ambient temperatures to fine-tune the depth of primary dormancy, should flowering happen earlier or later in the season. This is especially true in northern Sweden, where plants vernalize before winter (Duncan *et al.*, 2015) and usually flower as soon as the snow melts (day length and temperature permitting), the timing of which is likely to vary from year to year.

On the other hand, it is difficult to make sense of the great variability in the response to low temperatures observed among northern lines (Fig. 2 and 3B), as one would expect low dormancy levels to be necessary to make the most of an extremely short growing season (which is the norm at these latitudes). This suggest that these lines, despite their common geographical origin and high vernalization requirement (Shindo *et al.*, 2006; Duncan *et al.*, 2015), have different germination phenologies. Alternatively, although modelling approaches predict that reproduction occurs under similar temperatures across the species range (Springthorpe and Penfield, 2015), it is possible that populations from northern Sweden set seeds in slightly warmer temperatures (Burghardt *et al.*, 2016b), which would diminish the environmental effect and result in weaker dormancy.

Germination in Sweden also happens — to a much lesser extent — in spring and/or summer, as it is not uncommon to observe flowering plants at different times of the year in some southern Swedish populations (Kerdaffrec and Nordborg, personal observations). This could be evidence of bet-hedging, and it is clear that, in this case, the ability to adjust dormancy levels through maternal regulation according to seasonal environment would be advantageous. However, a constant monitoring of these populations over several years would be necessary to rule out the possibility that distinct genotypes expressing different life cycles segregate at these locations.

### The genetic basis of the response to low seed maturation temperatures

Although the molecular mechanisms involved in the temperature-dependent maternal regulation of seed dormancy are being revealed, its underlying genetic basis has not been studied yet. Here, by performing a GWAS on six dormancy traits, we identify a total of nine distinct associations, three of which have a ‘specific’ (i.e., interaction) effect (Table 3). SNPs within these three peaks are associated with high germination rates in response to cold seed maturation temperatures, which suggests that they tag genes involved in the temperature-dependent regulation of dormancy.

Among the candidates for the ‘specific’ genes, we identified *PHOT1* and *PHOT2,* which both encode phototropins that mediate several light-dependent processes such as hypocotyl phototropism (Zhao *et al.*, 2013), stomatal opening (Kinoshita *et al.*, 2001) and germination (Jedynak *et al.*, 2013). Light, along with temperature, is one of the factors regulating primary dormancy induction, and later in the soil seed bank dormancy release and germination. Light-induced germination is mainly promoted by phytochromes, especially PHYB and PHYA (Shinomura *et al.*, 1994; Heschel *et al.*, 2007; Jiang *et al.*, 2016), and phototropins are assumed to act downstream of them, by modulating the germination response via the integration of light and temperature signals (Jedynak *et al.*, 2013). Interestingly, *TRANSPARENT TESTA 12* (*TT12*), a gene central to the induction of coat-imposed dormancy in response to low seed maturation temperatures (MacGregor *et al.*, 2015), was identified earlier in a GWAS for germination traits under various light treatments (Morrison and Linder, 2014). This stresses the point that light and temperature signaling pathways may interact both during the induction and the release of dormancy. Therefore, it is possible that *PHOT1* and *PHOT2* play a role in dormancy regulation, direct evidence of which remains to be established. Finally, it should be mentioned that *TT12* and *PHYTOCHROME-INTERACTING FACTOR-LIKE 6* (*PIL6*), a gene negatively regulating *PHYB* (Fujimori *et al.*, 2004), are located 45 kb and 35 kb downstream of the strongest association detected in our analysis (peak 6, chr.3), respectively.

Although our GWAS identified compelling candidates, we emphasize that the signals are driven by few lines, a consequence of the small sample size and the limited phenotypic variation. Indeed, most of the associated SNPs, and especially those with a ‘specific’ effect, are relatively rare and often private to a small subset of non-dormant, cold-temperature-insensitive northern lines (Table 3). Some of these associations may also be false positives due to confounding by population structure, although quantile-quantile plots do not show extremely inflated *P* values (Fig. S5).

As previously mentioned, both the power and the resolution of our GWAS are undermined by the limited dormancy variation observed among Swedish lines. Conspicuously, almost half of the tested lines are deeply dormant (‘noninformative’ lines). In future experiments, it could be interesting to focus only on lines with mild- or non-dormant phenotypes, or alternatively, to apply variable cold stratification treatments to gradually alleviate dormancy and maximize the variation. On the other hand, classical quantitative trait locus (QTL) mapping could be performed in segregating populations derived from contrasted lines such as, for example, Gro-3 (insensitive) and Löv-1 (sensitive) (Fig. S2).

Finally, we have previously shown by performing a GWAS on 161 Swedish lines that *DOG1* is the major regulator of seed dormancy in Sweden (Kerdaffrec *et al.*, 2016). The *DOG1* region was also associated in the present study (Fig. S4), but to a lesser degree, although similar phenotypes were used in both cases (GR21 warm). There are two likely reasons for this discrepancy. First, the phenotypes are not perfectly correlated (*r* = 0.84) and several lines were slightly more dormant in this study than in the previous (Fig. S6), reflecting the plastic nature of seed dormancy. Secondly, even if the previously identified *DOG1* alleles segregate among the lines used here, a different sample size is likely to give different results because of the pitfalls intrinsic to GWAS (altered power, changes in allele frequencies, epistasis, among others) (Korte and Farlow, 2013). As a demonstration, a GWAS on both GR21 warm phenotypes (previous and present) using the exact same set of lines (86, the overlap between both studies) gave very similar results at the *DOG1* locus (Fig. S7).

### Towards a better understanding of environmentally-dependent life-history transitions

In this study, we confirm that the maternal environment interacts with genotype in controlling seed dormancy variation and characterize this interaction in a natural variation context. Our GWAS results, in spite of their limitations, clearly support the fact the maternal environment impacts the genetic basis of seed dormancy.

Because the gene networks and signaling pathways involved in the regulation of environmentally-dependent transitions are starting to be well characterized, it will become increasingly possible to integrate them into predictive models that can later be validated in field experiments. The need for a better understanding of the molecular genetic basis of genotype-environment interactions is real, as they are of importance not only to evolutionary biology (Via and Lande, 1985; Fournier-Level *et al.*, 2011), but also to modern agriculture, especially in the light of climate change (Saranga *et al.*, 2001; Li *et al.*, 2014; Fournier-Level *et al.*, 2016).

## Supplementary material

Table S1. List of Swedish lines used in this study.

Table S2. List of *a priori* seed dormancy genes used in this study.

Table S3. Top associated *a priori* seed dormancy genes.

Figure S1. Pairwise correlations between the dormancy traits and latitude.

Figure S2. The diverse germination trajectories in Sweden.

Figure S3. Enlarged view of the region surrounding peak 9 (1 Mb).

Figure S4. Enlarged view of the region surrounding peak 8 (1 Mb).

Figure S5. Quantile-quantile plots of GWAS *P* values.

Figure S6. The relationship between present and previously published GR21 phenotypes.

Figure S7. *DOG1* region association scans for GR21 warm phenotypes.

## Acknowledgments

This work was supported by European Research Council grant 268962 (MAXMAP) to MN. We thank Eriko Sasaki for help with LIMIX, Danièle L Filiault for constructive feedback regarding analyses and manuscript, and other members of the Nordborg lab for useful discussions.

